# Receptor-binding domain 2 of *Clostridioides difficile* binary toxin as a promising vaccine component against *C. difficile* infection

**DOI:** 10.64898/2026.08.30.748174

**Authors:** Shaohui Wang, Joshua Heuler, Yukihiro Nakanishi, Hyeun Bum Kim, Xingmin Sun

## Abstract

Symptoms of *Clostridioides difficile infection* (CDI) are primarily caused by two major protein toxins, toxin A (TcdA) and toxin B (TcdB). In addition, approximately 5-30% of *C. difficile* strains produce a third toxin, *C. difficile* binary toxin (CDT), which is has been associated with enhanced virulence and severe disease. CDT consists of an enzymatic component CDTa, and a binding and translocation component CDTb, which mediates the delivery of CDTa into host cells. CDTb contains two receptor-binding domains, RBD1 and RBD2. Recent structural studies suggest that RBD2 plays a critical role in the formation and stabilization of the di-heptameric CDTb assembly required for efficient intoxication of host cells. In this study, we evaluated the immunogenicity and protective potential of RBD1 and RBD2 using *in silico, in vitro* and *in vivo* approaches. Sequence analysis demonstrated that RBD2 is highly conserved among diverse CDT-producing *C. difficile* ribotypes and toxinotypes. Immunization of mice with RBD2, but not RBD1 conferred effective protection against direct CDT challenge. Moreover, RBD2 immunization protected hamsters against infection with a CDT-only-producing *C. difficile* strain (DSM 101085; TcdA⁻TcdB⁻CDT⁺). Mechanistically, anti-RBD2 serum, but not anti-RBD1 serum, effectively neutralized CDT-mediated cytotoxicity, as demonstrated by inhibition of cell rounding in Vero cells. Collectively, these findings identify RBD2 as a promising vaccine antigen targeting CDT and provide functional evidence supporting its critical role in CDT-mediated host-cell intoxication. Incorporation of RBD2 into multivalent *C. difficile* vaccines may broaden protection against hypervirulent, CDT-producing strains.

## Introduction

*Clostridioides difficile* is an anaerobic, spore-forming, Gram-positive bacterium ^1^. Symptoms of *C. difficile* infection (CDI) range from diarrhea and intestinal inflammation to death and are mainly caused by two protein toxins, toxin A (TcdA) and toxin B (TcdB) ^2,3^. In addition to TcdA and TcdB, approximately 5-30% of *C. difficile* strains produce a third toxin termed binary toxin (CDT), which is encoded by the *cdtA* and *cdtB* genes^4,5^. CDT is believed to enhance TcdA- and TcdB-mediated toxicity and is associated with severe disease and higher recurrence rates ^6,7^. Strains producing only the CDT (A-B-CDT+) retain virulence in the clinic ^8,9^. Despite a well-acknowledged clinical need and more than two decades extensive efforts from both academia and pharmaceutical industry, no vaccine against CDI has been licensed ^10–12^. Previous clinical vaccine candidates (e.g., VLA84, Sanofi, Pfizer) have targeted only TcdA and TcdB. However, a fully effective vaccine may need to target all three toxins.

CDT is composed of an enzymatic component CDTa, and a binding and translocation component, CDTb, which mediates the cell entry of CDTa into host cells ^13^. CDTa is a two-domain enzyme: the N-terminal region (residues 1 to 215), which is important for CDTa binding to CDTb, and the C-terminal region (residues 224 to 420), which is important for enzyme’s catalytic toxicity and catalyzes the ADP-ribosylation of actin ^14,15^. CDTb is the binding and pore-forming component of the CDT. CDTb is synthesized as an inactive precursor form including SD (signaling domain), AD (activation domain), HD1 (heptamerization domain 1), βBD (β binding domain), HD2 (heptamerization domain 2), HD3 (heptamerization domain 3), RBD1 (receptor binding domain 1) and RBD2 (receptor binding domain 2) ^16,17^. The precursor form CDTb undergoes proteolytic cleavage, with removal of SD and AD, to generate mature and activated CDTb (**mCDTb**)^18^. Removal of SD and AD can be achieved by incubation of precursor CDTb with trypsin *in vitro*.

Recent structural studies of CDTb showed that RBD1 domain lacks sequence homology to any other known toxin and contains a Ca^2+^-binding site, whereas the RBD2 domain is not present in other members of this toxin family. Also, RBD2 was also shown to be critical for establishing the di-heptamer macromolecular assembly in the activated CDTb that is necessary for host cell toxicity ^18^. Deletion of RBD2 from CDTb, resulted in a heptameric structure with significantly reduced cytotoxicity. It was postulated that the RBD2 domain is crucial for the di-heptamer assembly and is also critical for delivering toxic CDTa to host cells ^14^. These studies provide a structural basis for mapping RBD2-mediated molecular interactions and for developing inhibitors targeting CDTb.

In this study, we analyzed the cytotoxic effect of CDTa/b, evaluated the immunogenicity of RBD1 and RBD2 and their potential as effective vaccine components against CDI.

## Results

### Homology of CDTb and RBD2 among major toxinotypes and ribotypes of *C. difficile* strains

An effective vaccine candidate should be highly conserved across different ribotypes (RTs) of *C. difficile*. To this end, we investigated the sequence conservation of CDTb and RBD2 among major CDT-producing *C. difficile* toxinotypes (TcdA⁺TcdB⁺CDT⁺, TcdA⁻TcdB⁺CDT⁺, and TcdA⁻TcdB⁻CDT⁺) and RTs (RT027, RT078, RT066, RT126, RT045, RT033, RT023, RT019, RT036, and RT244) (Table 1). A maximum-likelihood phylogenetic tree was generated using CDTb amino acid sequences from representative strains of CDT-positive RTs (Fig. 1). The results suggest a degree of correlation between RT and CDTb sequence similarity. CDTb sequences from RT023 were identical and formed a distinct cluster in the phylogenetic tree, separate from sequences derived from other ribotypes. Similar patterns were observed for CDTb sequences from RT019 and RT036. In some cases, multiple RTs shared nearly identical CDTb sequences. For example, CDTb sequences from representative strains of RT033, RT045, RT066, RT078, and RT126 were virtually identical. CDTb sequences from RT027 and RT244 also clustered closely together. Overall, CDTb sequences from strains belonging to the same RT clustered together, suggesting that CDTb is highly conserved at the amino acid level within individual RTs. However, because only two to three representative sequences were analyzed for each RT, a larger-scale analysis may reveal exceptions to this trend.

**Fig. 1.**
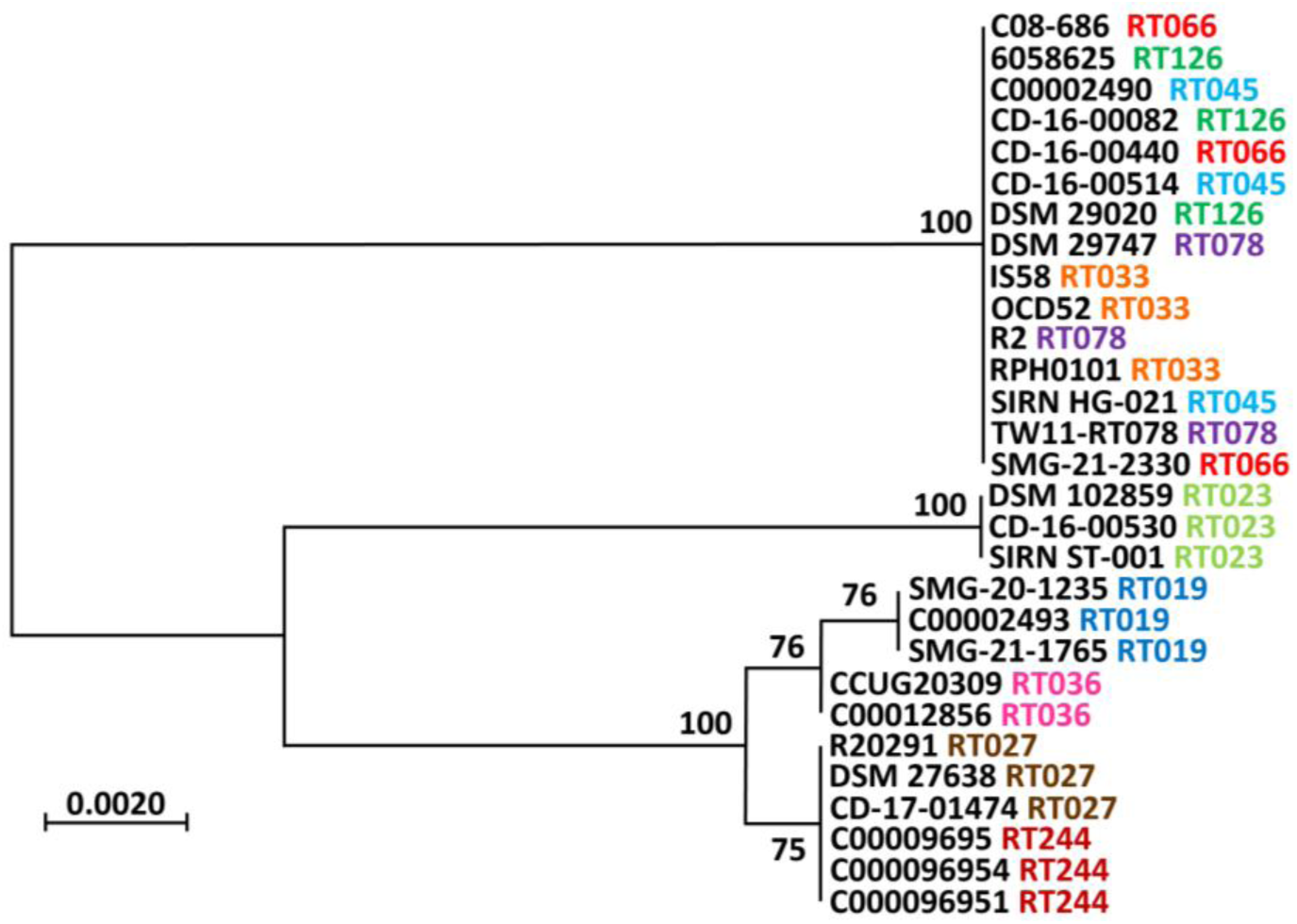
Phylogenetic analysis of CDTb amino acid sequences. CDTb amino acid sequences were aligned using the MUSCLE algorithm in MEGA X, and a maximum-likelihood phylogenetic tree was constructed with 500 bootstrap replicates. Bootstrap values >50 are displayed at the corresponding nodes. The ribotype (RT) of each source strain is indicated adjacent to the strain name and color-coded for ease of identification.

**Table 1.**
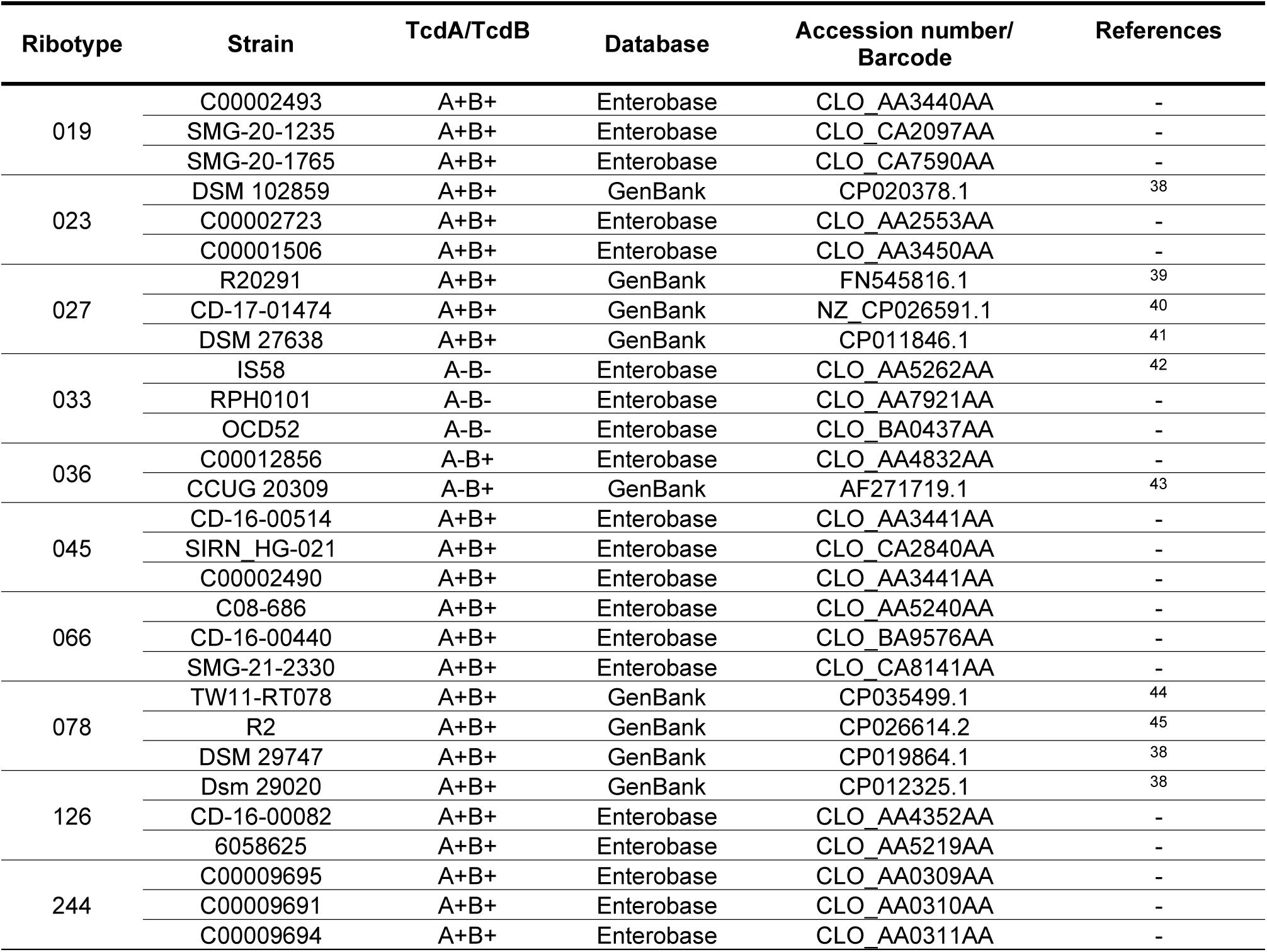
*C. difficile* strains selected for CDTb analysis.

To further examine CDTb sequence diversity, we aligned the full-length CDTb (Fig. 2) and RBD2 domain (Supplemental Fig. 1) amino acid sequences using MUSCLE and visualized the alignments with Jalview. Domains were annotated based on a previous review^19^. Sequence variations were most frequent in the signaling domain (1 variation per 7.33 residues), followed by the activation domain (1 variation per 12.92 residues), RBD2 (1 variation per 20 residues), RBD1 (1 variation per 25.8 residues), and heptamerization domain 3 (1 variation per 34.33 residues). In contrast, heptamerization domain 1, heptamerization domain 2, and the β-barrel domain were completely conserved among the strains examined. Within RBD2 specifically, the six identified sequence variations were concentrated toward the N-terminal region of the domain (residues 762–789), whereas the C-terminal region was highly conserved. Collectively, these analyses demonstrate that both CDTb and RBD2 are highly conserved among the *C. difficile* strains examined.

**Fig. 2.**
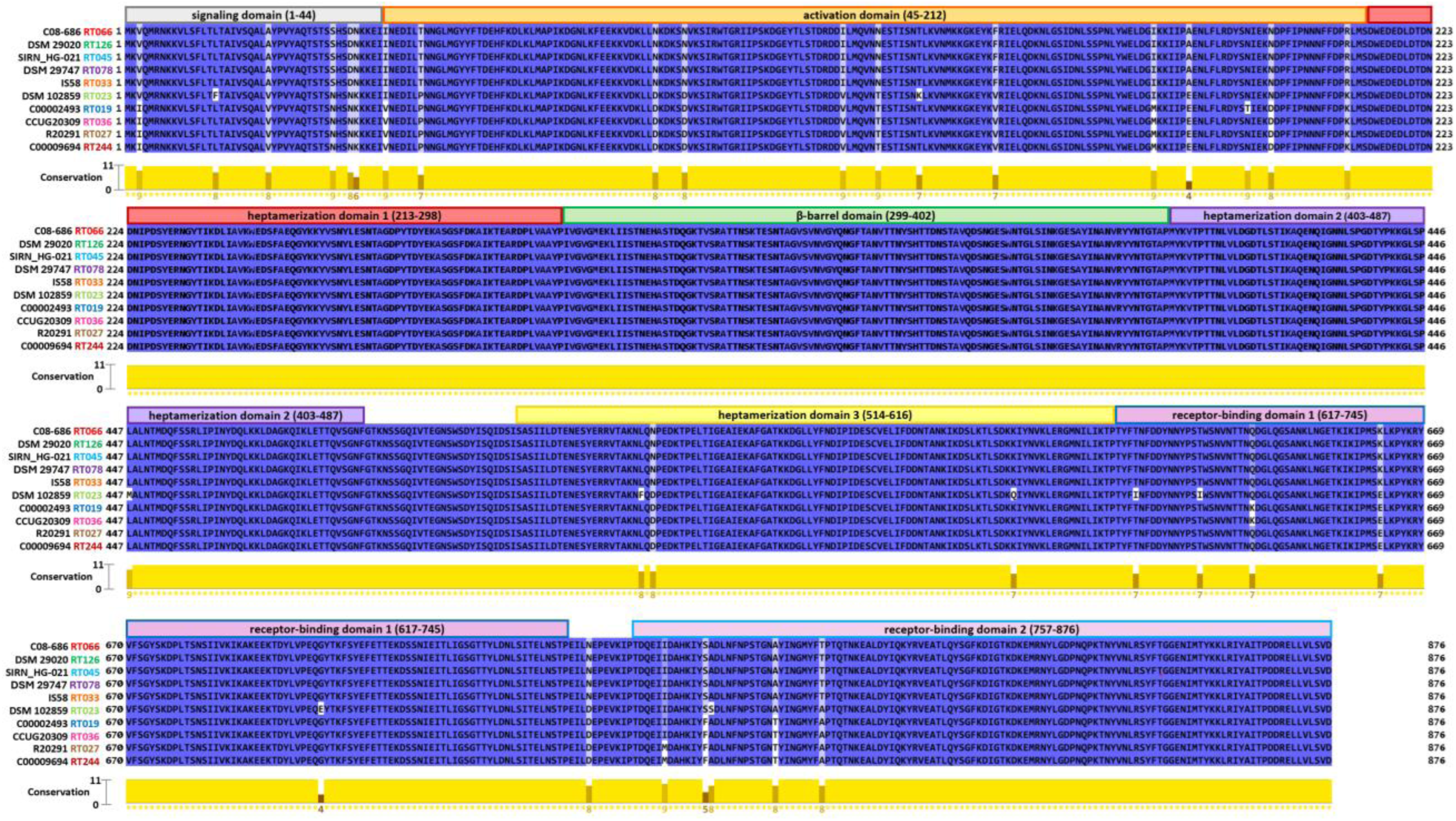
Sequence conservation of CDTb domains among representative *C. difficile* strains. MUSCLE alignments of CDTb amino acid sequences were visualized using Jalview. Conservation scores ranging from 0 (no conservation) to 11 (complete conservation) were calculated for each amino acid position using Jalview (see Methods). The ribotype (RT) of each source strain is indicated adjacent to the strain name and color-coded for ease of identification.

### Both RBD1 and RBD2 contain abundant predicted B-cell epitopes

To assess the potential immunogenicity of RBD1 and RBD2, we performed B-cell epitope prediction using the BepiPred 2.0 server (https://www.iedb.org/). The analysis revealed that both RBD1 and RBD2 contain multiple regions predicted to represent linear B-cell epitopes (Fig. 3, highlighted in yellow), suggesting that both domains have substantial potential to elicit antibody responses.

**Fig. 3.**
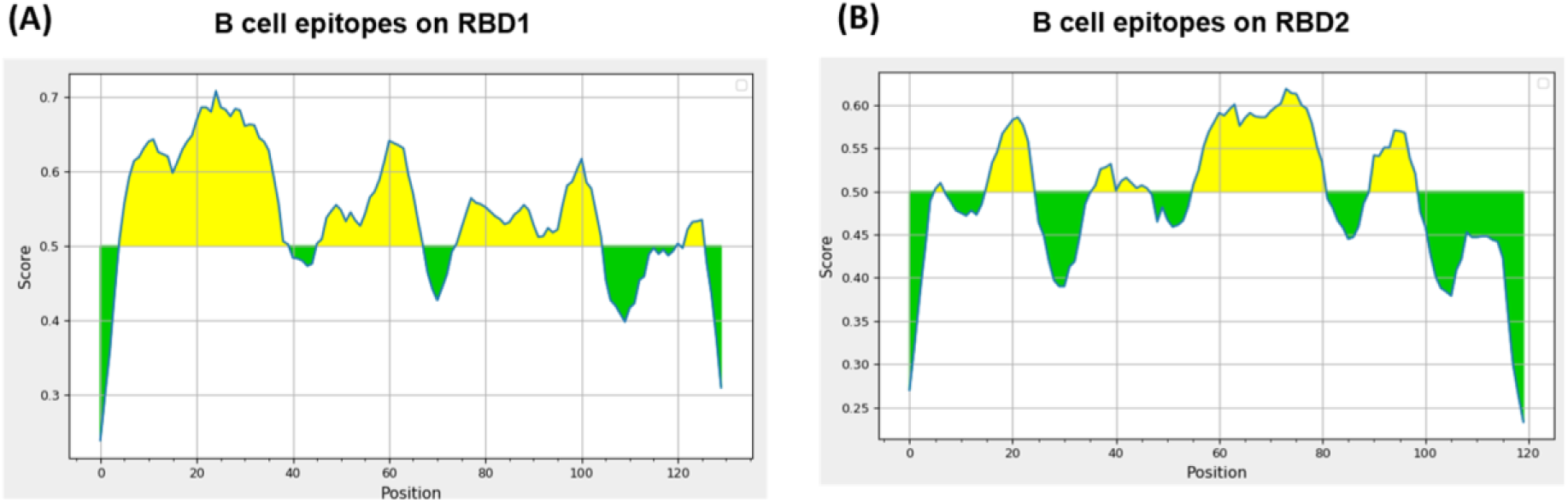
Predicted linear B-cell epitopes within RBD1 and RBD2. Linear B-cell epitopes within RBD1 and RBD2 were predicted using the BepiPred-2.0 server (https://www.iedb.org/). Amino acid residues with prediction scores above the default threshold of 0.5 were considered potential B-cell epitope residues and are highlighted in yellow. The y-axis represents the BepiPred-2.0 prediction score, and the x-axis represents the amino acid position within the protein sequence.

### CDT (CDTa:mCDTb at 1:7 molar ratio) causes cell rounding in a concentration-dependent manner

To evaluate the immunogenicity of RBD1 and RBD2, we expressed and purified 6×His-tagged RBD1, RBD2, and RBD1+RBD2 using the *E. coli* BL21 expression system (Fig. 4C). CDTa and CDTb containing a 6×His tag were also expressed in *E. coli* BL21(DE3) and purified by Ni-affinity chromatography to >95% purity for subsequent evaluation of CDT cytotoxicity and antibody-mediated neutralization (Fig. 4B).

**Fig. 4.**
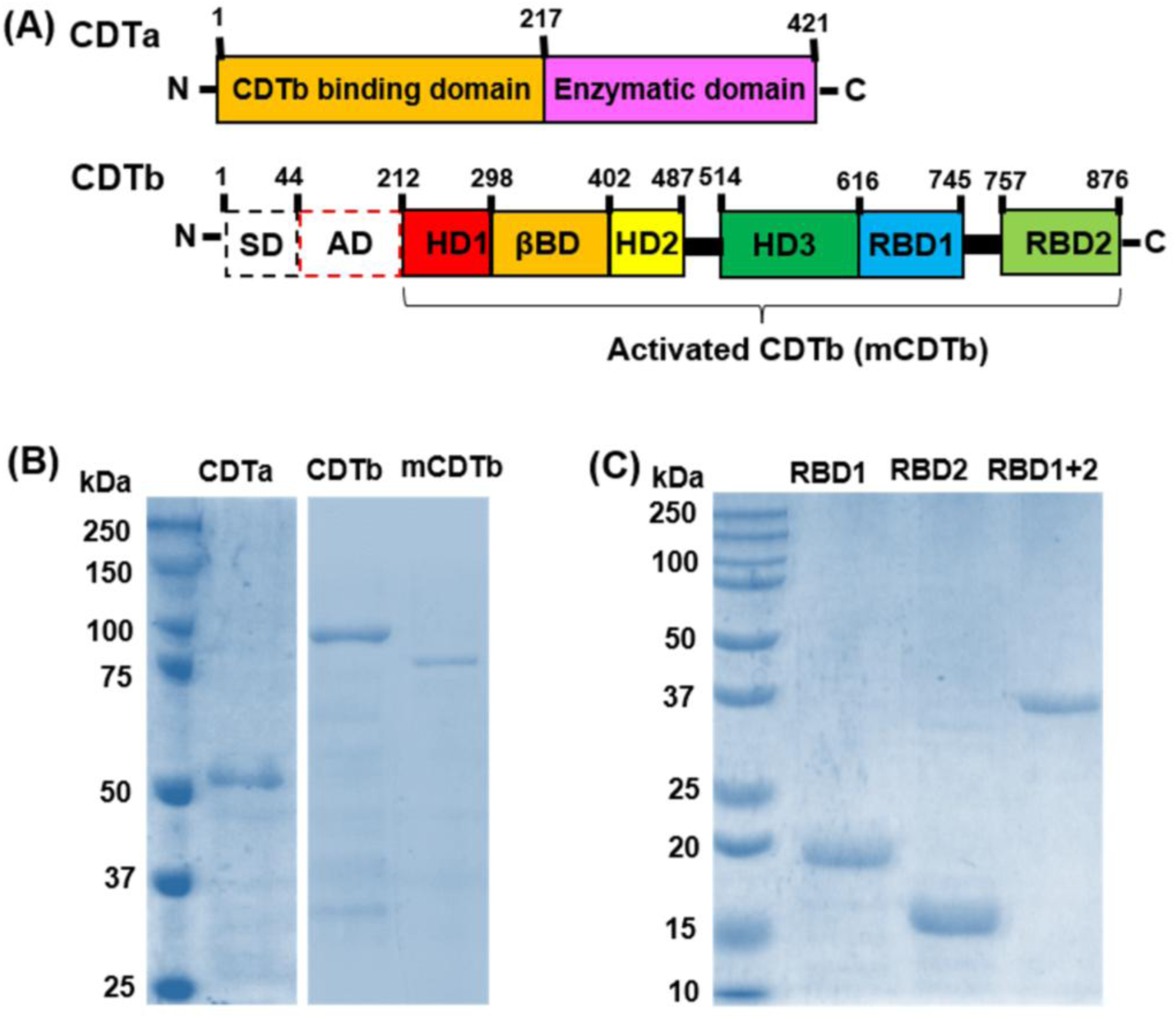
Domain organization, expression, purification, and activation of CDT components and CDTb receptor-binding domains. **(A)** Domain organization of CDTa and CDTb. CDTb is synthesized as an inactive precursor containing SD (signaling domain), AD (activation domain), HD1 (heptamerization domain 1), βBD (β-binding domain), HD2 (heptamerization domain 2), HD3 (heptamerization domain 3), RBD1 (receptor-binding domain 1), and RBD2 (receptor-binding domain 2). Proteolytic removal of SD and AD generates mature, activated CDTb (mCDTb). *In vitro*, CDTb activation was achieved by trypsin treatment. **(B)** Expression, purification, and activation of CDTa and CDTb. Recombinant CDTa and CDTb were expressed in E. coli, purified from bacterial lysates by Ni-affinity chromatography, and analyzed by SDS-PAGE. CDTb was activated by incubation with trypsin (0.2 μg trypsin/μg protein) for 30 min at 37°C to generate mCDTb. **(C)** Expression and purification of RBD1, RBD2, and RBD1+RBD2 (RBD1+2). Recombinant RBD1, RBD2, and RBD1+2 were expressed in *E. coli*, purified from bacterial lysates by Ni-affinity chromatography, and analyzed by SDS-PAGE.

Activated CDTb (mCDTb) was generated by incubating CDTb with trypsin at 0.2 μg of trypsin/μg of protein for 30 min at 37°C (Fig. 4B). The cytotoxicity of CDTa, CDTb, mCDTb, or their combinations was evaluated using Vero cells. CDTa, CDTb, or mCDTb alone did not induce cell rounding at any of the concentrations tested (Fig. 5A). In contrast, the combination of CDTa and mCDTb at concentrations of 100 ng/mL CDTa and 1,104 ng/mL mCDTb (CDTa at a 1:7 molar ratio), or at higher concentrations, induced the characteristic rounding of Vero cells after 2 h of incubation (Fig. 5). Interestingly, both RBD2 and RBD1+RBD2, but not RBD1, significantly inhibited CDT-mediated cell rounding (Fig. 6), suggesting an essential role for RBD2 in CDT binding to Vero cells and subsequent CDT-mediated cytotoxicity.

**Fig. 5.**
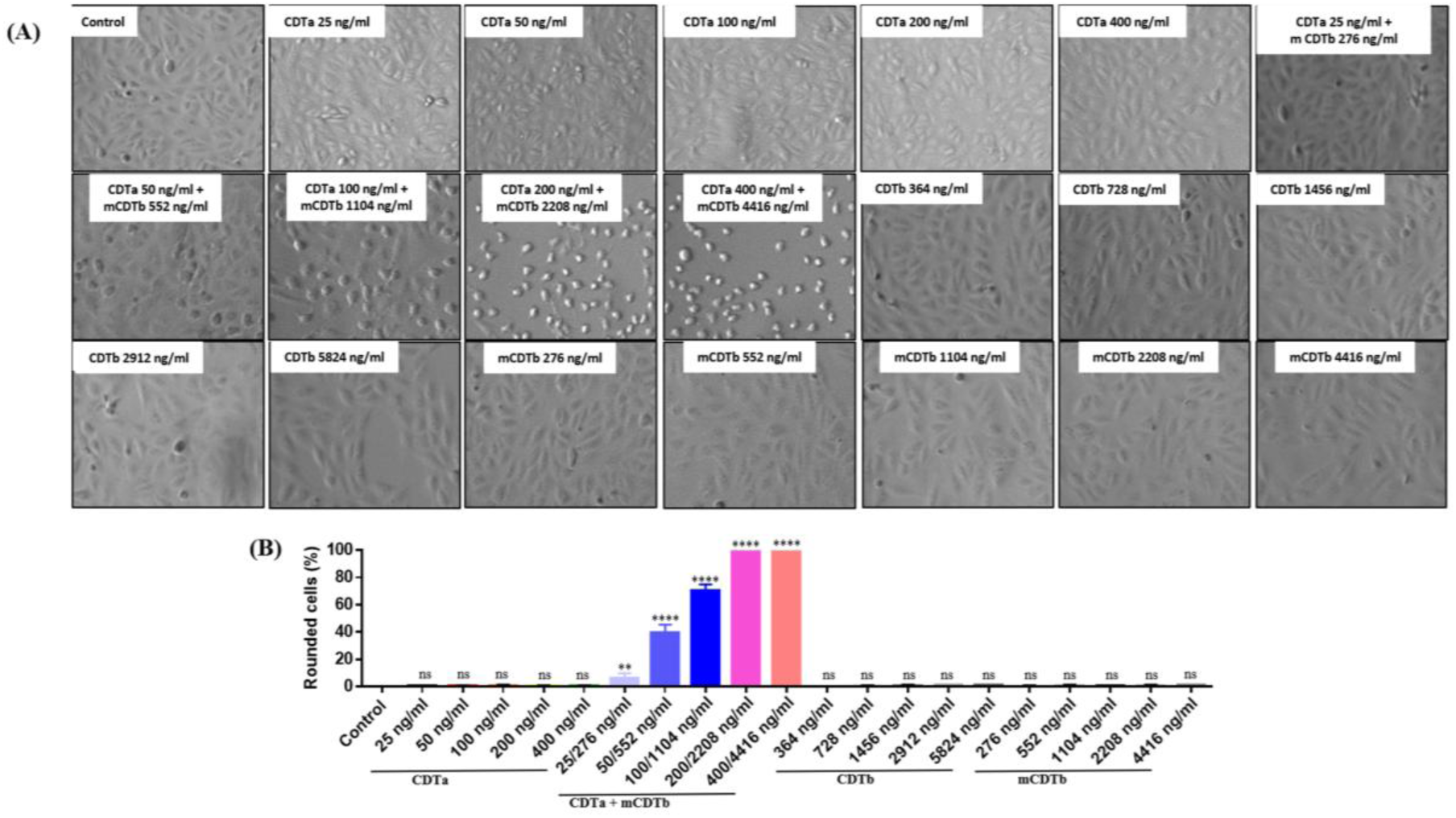
CDT induces concentration-dependent cell rounding in Vero cells. Vero cells were treated with increasing concentrations of CDTa, CDTb, or activated CDTb (mCDTb) alone, or with combinations of CDTa and mCDTb at the indicated concentrations and ratios. Untreated cells served as controls. **(A)** Representative images of Vero cells acquired after 2 h of treatment. **(B)** Quantification of the percentage of rounded cells. Data is presented as the mean ± SEM (n = 3 images per condition). Results from one representative experiment are shown. Statistical significance was determined by one-way ANOVA followed by Dunnett’s multiple-comparisons test. **p ≤ 0.01; ****p ≤ 0.0001; ns, not significant compared with the untreated control.

**Fig. 6.**
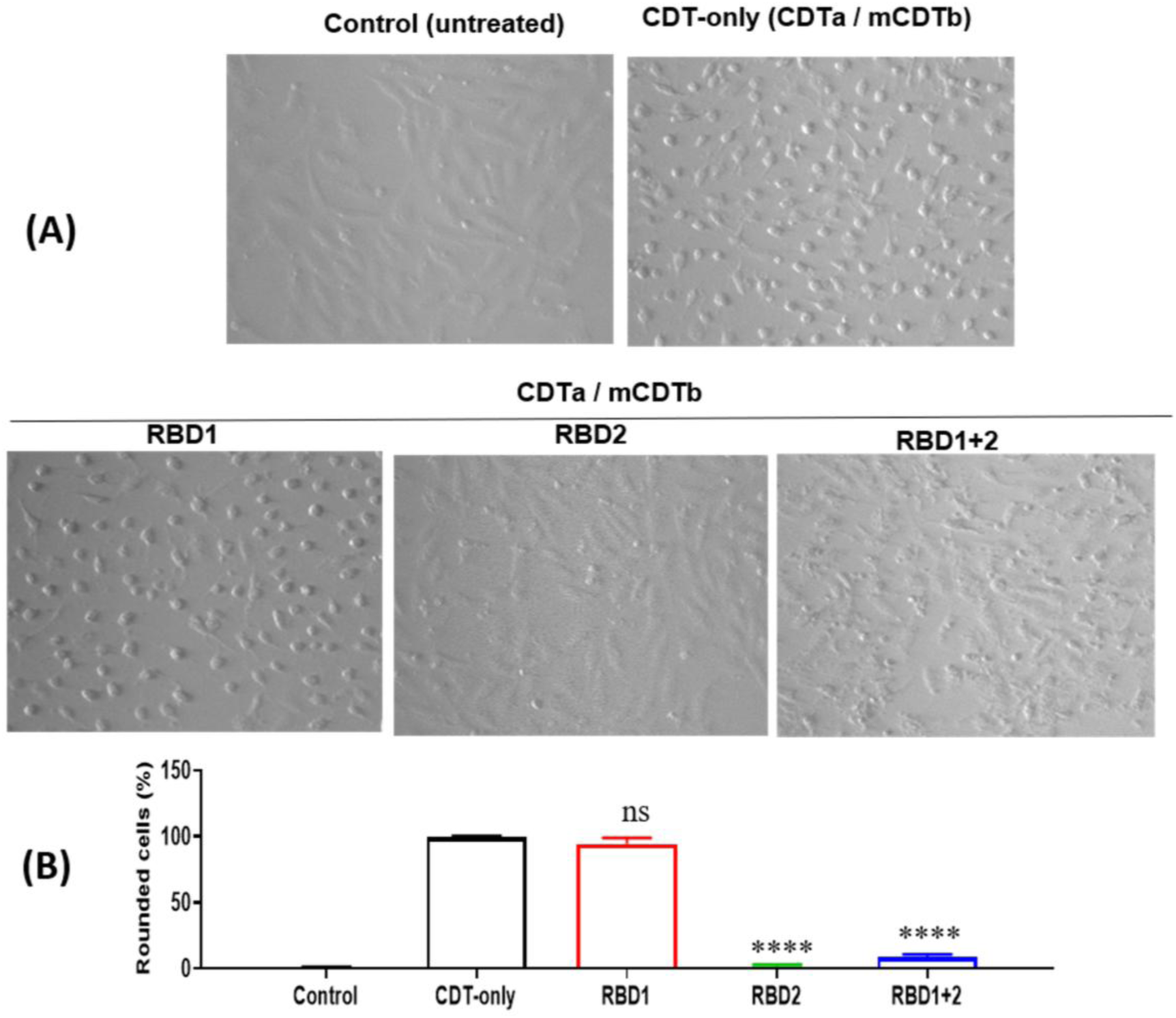
RBD2 and RBD1+RBD2 inhibit CDT-mediated cell rounding in Vero cells. **(A)** Vero cells were preincubated with RBD1, RBD2, or RBD1+RBD2 (1 µg/mL) for 30 min, followed by treatment with CDT consisting of 200 ng/mL CDTa and 2,208 ng/mL mCDTb (CDTa at a 1:7 molar ratio). Cells treated with CDT in the absence of RBD proteins served as the CDT-only control. Representative images were acquired after 2 h of CDT treatment. **(B)** The percentage of rounded cells was quantified from the acquired images. Data are presented as the mean ± SEM (n = 3 images per condition). Experiments were repeated three times, and results from one representative experiment are shown. Statistical significance was determined using Student’s t-test. ****p ≤ 0.0001; ns, not significant compared with the CDT-only control.

### Immunization with RBD2 or RBD1+2 induces significant antibody responses and protects mice against CDT challenge

With biologically active CDT established, the immunogenicity and protective efficacy of RBD1, RBD2, and RBD1+RBD2 were evaluated *in vitro* and *in vivo*. Intramuscular immunization of mice with 10 µg of RBD2 or 20 µg of RBD1+RBD2 adjuvanted with alum induced significant IgG and IgA antibody responses against RBD2 (Fig. 7) and CDTb (Fig. 8) in serum and fecal samples. Immunization with RBD1 using the same antigen dose and adjuvant induced significant anti-RBD1 IgG and IgA responses in serum (Fig. 7A, B) and a significant anti-RBD1 IgG response in fecal samples (Fig. 7C). However, RBD1 immunization did not induce a significant anti-RBD1 IgA response in fecal samples (Fig. 7D) or significant anti-CDTb IgG or IgA responses in either serum or fecal samples (Fig. 8). These results suggest that RBD2 is more immunogenic than RBD1 and induces antibody responses that more effectively recognize CDTb.

**Fig. 7.**
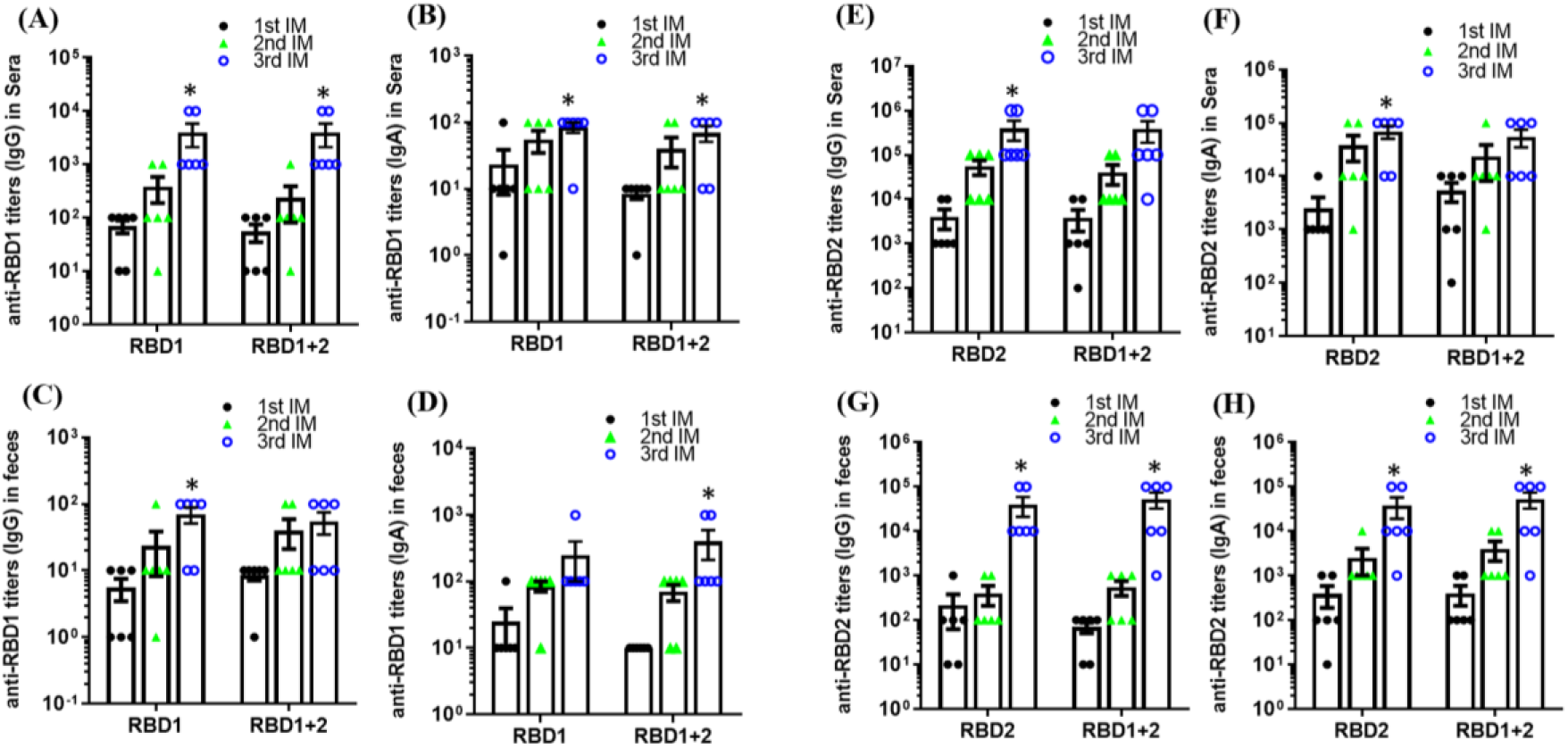
Intramuscular immunization with RBD1, RBD2, or RBD1+RBD2 induces significant antibody responses in mice. Groups of C57BL/6 mice (n = 6 per group) were immunized intramuscularly three times at 14-day intervals with 10 μg of RBD1, 10 μg of RBD2, or 20 μg of RBD1+RBD2, adjuvanted with alum. Serum and fecal samples were collected, and antigen-specific IgG and IgA titers were determined by ELISA. **(A–D)** Anti-RBD1 IgG and IgA antibody responses in serum and fecal samples. **(E–H)** Anti-RBD2 IgG and IgA antibody responses in serum and fecal samples. The experiment was independently performed twice, and data from one representative experiment are presented as the mean ± SEM (n = 6 per group). *p* < 0.05 compared with the first immunization (1st IM).

**Fig. 8.**
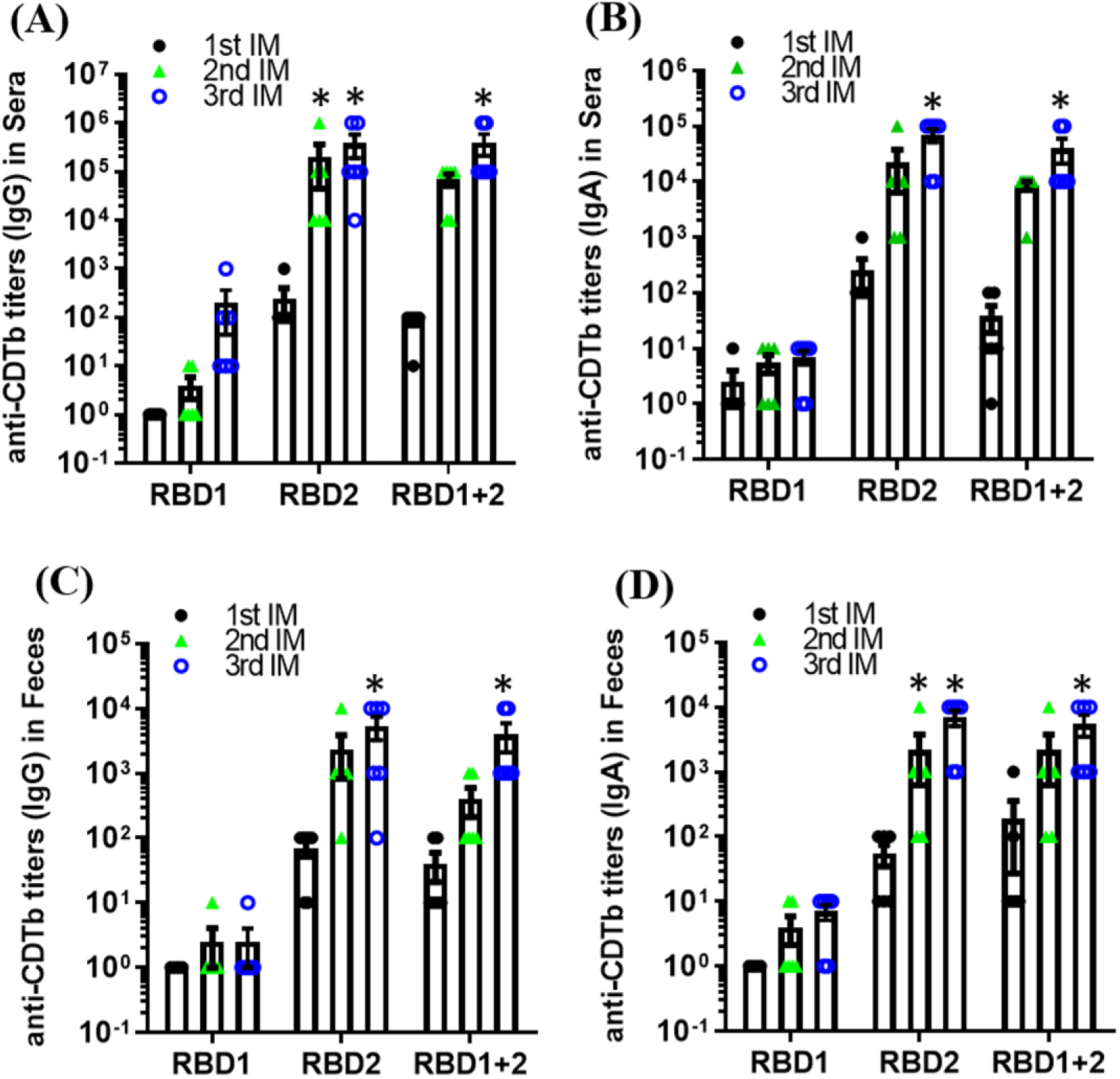
Intramuscular immunization with RBD2 or RBD1+RBD2 induces significant anti-CDTb antibody responses in mice. Groups of C57BL/6 mice (n = 6 per group) were immunized intramuscularly three times at 14-day intervals with 10 μg of RBD1, 10 μg of RBD2, or 20 μg of RBD1+RBD2, adjuvanted with alum. Serum and fecal samples were collected, and anti-CDTb IgG and IgA titers were determined by ELISA. **(A, B)** Anti-CDTb IgG and IgA responses in serum. **(C, D)** Anti-CDTb IgG and IgA responses in fecal samples. The experiment was independently performed twice, and data from one representative experiment are presented as the mean ± SEM (n = 6 per group). *p* < 0.05 compared with the first immunization (1st IM).

The protective efficacy of immunization with RBD1, RBD2, or RBD1+RBD2 was subsequently evaluated in mice. After three immunizations, mice were challenged with a lethal dose of CDT consisting of 260 ng of CDTa and 2,870 ng of mCDTb. In the non-immunized and RBD1-immunized groups, all mice died within 20 h after CDT challenge (Fig. 9A). In contrast, all mice immunized with either RBD2 or RBD1+RBD2 survived the challenge (Fig. 9A). No deaths occurred among non-immunized or RBD1-, RBD2-, or RBD1+RBD2-immunized mice challenged with mCDTb alone (Fig. 9B), confirming that mCDTb alone was not lethal under the conditions tested. Collectively, these results demonstrate that RBD2 elicits stronger CDTb-reactive antibody responses than RBD1 and that immunization with RBD2, either alone or in combination with RBD1, provides complete protection against lethal CDT challenge in mice.

**Fig. 9.**
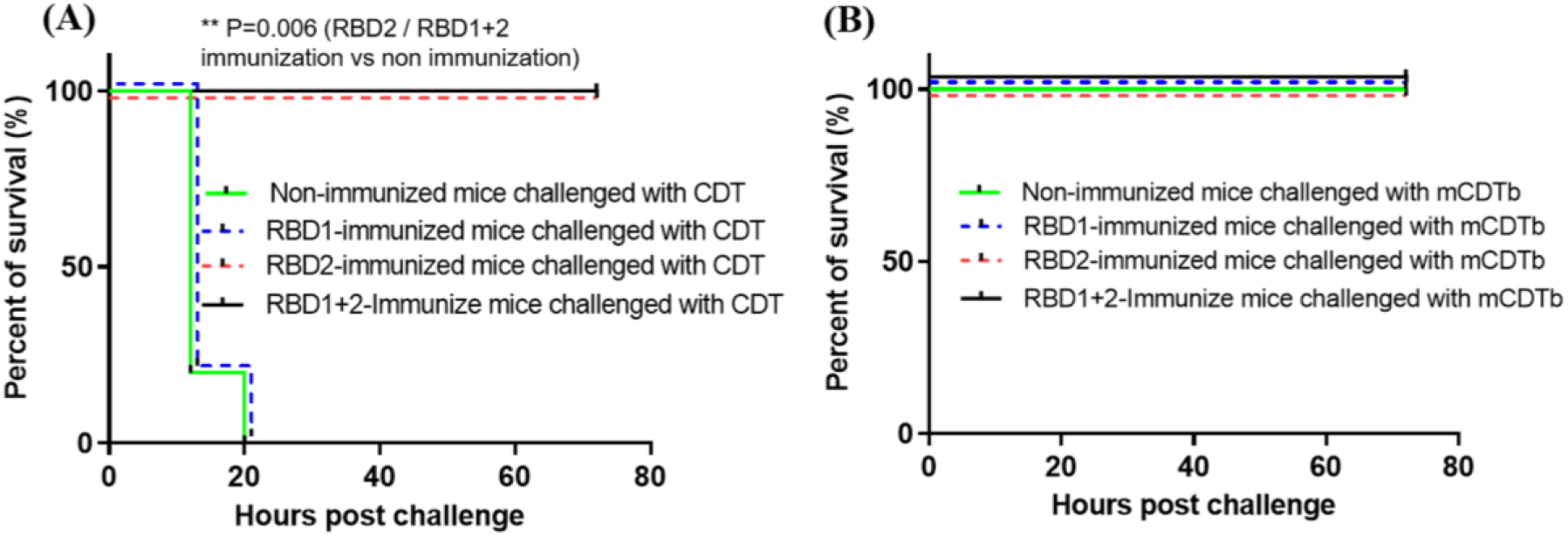
Immunization with RBD2 or RBD1+RBD2, but not RBD1, protects mice against lethal CDT challenge. (A) Fourteen days after the third immunization, mice immunized with RBD1, RBD2, or RBD1+RBD2, as well as non-immunized control mice (n = 6 per group), were challenged intraperitoneally (i.p.) with a lethal dose of CDT consisting of 260 ng of CDTa and 2,870 ng of mCDTb per mouse (CDTa at a 1:7 molar ratio). Mice were monitored for survival and clinical signs of disease for 80 h after challenge. (B) mCDTb alone is not lethal to mice. Fourteen days after the third immunization, mice immunized with RBD1, RBD2, or RBD1+RBD2, as well as non-immunized control mice (n = 6 per group), were challenged i.p. with mCDTb alone (2,870 ng/mouse) and monitored for survival and clinical signs of disease for 72 h. Survival differences were analyzed using the Kaplan–Meier method with the log-rank (Mantel–Cox) test. **P = 0.006 for RBD2- and RBD1+2-immunized mice compared with non-immunized control mice.

### Anti-RBD2 and anti-RBD1+RBD2 sera inhibit CDT-mediated cell rounding

We next investigated whether antibodies induced by RBD immunization could neutralize CDT-mediated cytotoxicity. Confluent Vero cells were treated with CDT (200 ng/mL CDTa plus 2,208 ng/mL mCDTb; CDTa at a 1:7 molar ratio) in the presence of pre-immune serum or anti-RBD1, anti-RBD2, or anti-RBD1+RBD2 serum for 2 h. As shown in Fig. 10, at a 1:400 serum dilution, anti-RBD2 and anti-RBD1+RBD2 sera significantly reduced CDT-mediated cell rounding to 45 ± 5% and 39.33 ± 6.67%, respectively, compared with 100% cell rounding in the absence of serum. At a 1:50 dilution, anti-RBD2 and anti-RBD1+RBD2 sera almost completely inhibited CDT-mediated cell rounding, whereas anti-RBD1 serum did not exhibit comparable inhibitory activity. These results demonstrate that antibodies elicited by RBD2, either alone or in combination with RBD1, effectively neutralize CDT-mediated cytotoxicity *in vitro*.

**Fig. 10.**
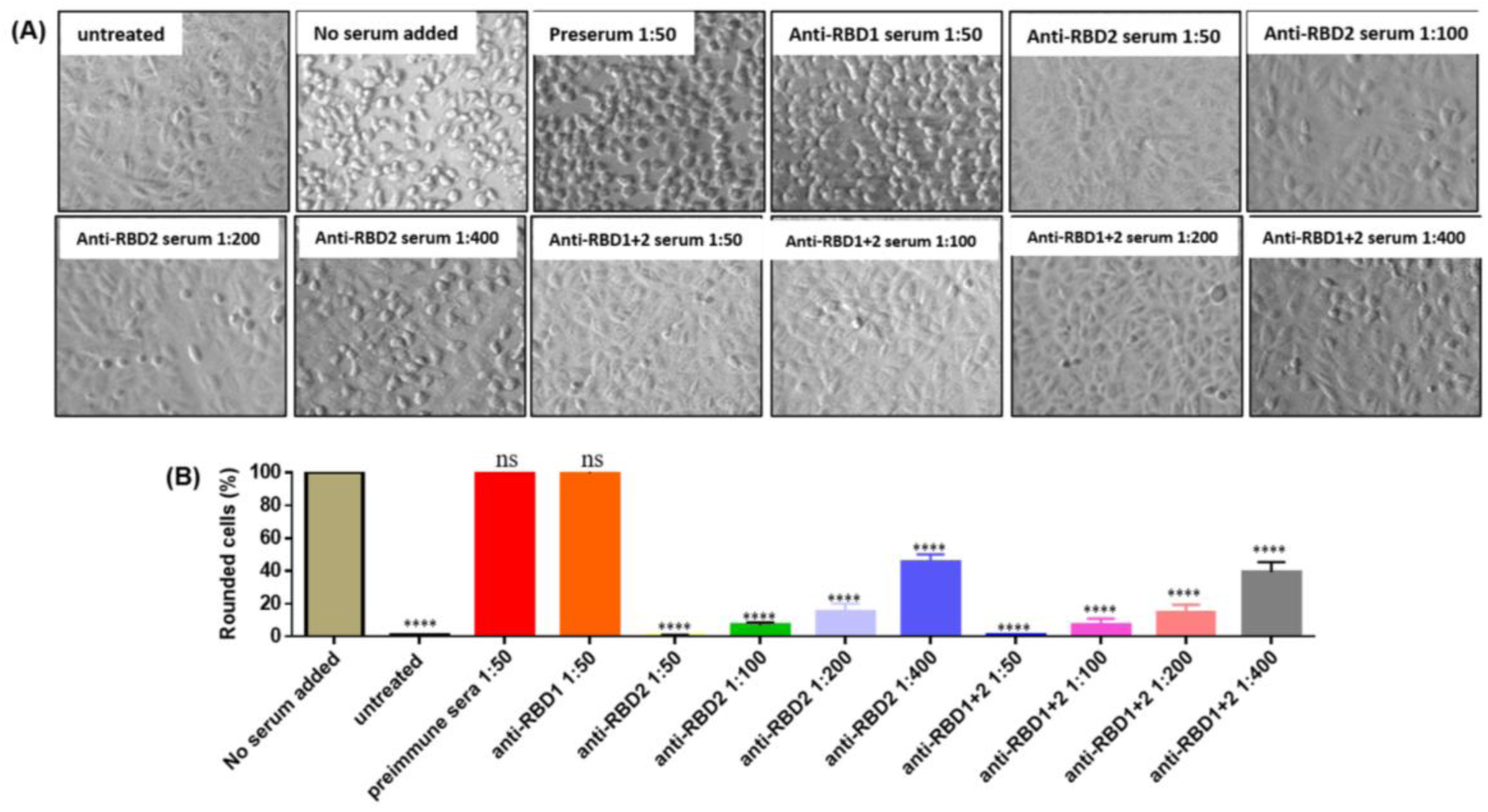
Anti-RBD2 and anti-RBD1+RBD2 sera inhibit CDT-mediated cell rounding. Vero cells were treated with CDT consisting of 200 ng/mL CDTa and 2,208 ng/mL mCDTb (CDTa at a 1:7 molar ratio) in the presence of pre-immune serum or anti-RBD1, anti-RBD2, or anti-RBD1+RBD2 serum at 37°C. (A) Representative images acquired after 2 h of treatment. (B) Quantification of the percentage of rounded cells. Data are presented as the mean ± SEM (n = 3 images per condition). Experiments were independently performed three times, and results from one representative experiment are shown. Statistical significance was determined using one-way ANOVA followed by Dunnett’s multiple-comparisons test. ****p ≤ 0.0001; ns, not significant compared with cells treated with CDT in the absence of serum.

### Immunization of hamsters with RBD2 protein induces protective responses against infection with *C. difficile* strain DSM101085

Because hamsters are highly susceptible to CDI, we further evaluated the immunogenicity and protective efficacy of RBD2 in hamsters against infection with the CDT-only-producing *C. difficile* strain DSM 101085. Hamsters were immunized intramuscularly with RBD2 (30 µg per hamster per immunization) three times at 14-day intervals. RBD2 immunization induced significant anti-RBD2 IgG and IgA antibody responses in serum and fecal samples (Fig. 11A-D). Moreover, RBD2 immunization provided significant protection against challenge with 2 × 10⁴ spores per hamster of *C. difficile* strain DSM 101085 (A⁻B⁻CDT⁺) (Fig. 11E, F). These results demonstrate that RBD2 is immunogenic in hamsters and that RBD2 immunization provides significant protection against infection with a *C. difficile* strain that produces CDT but lacks TcdA and TcdB.

**Fig. 11.**
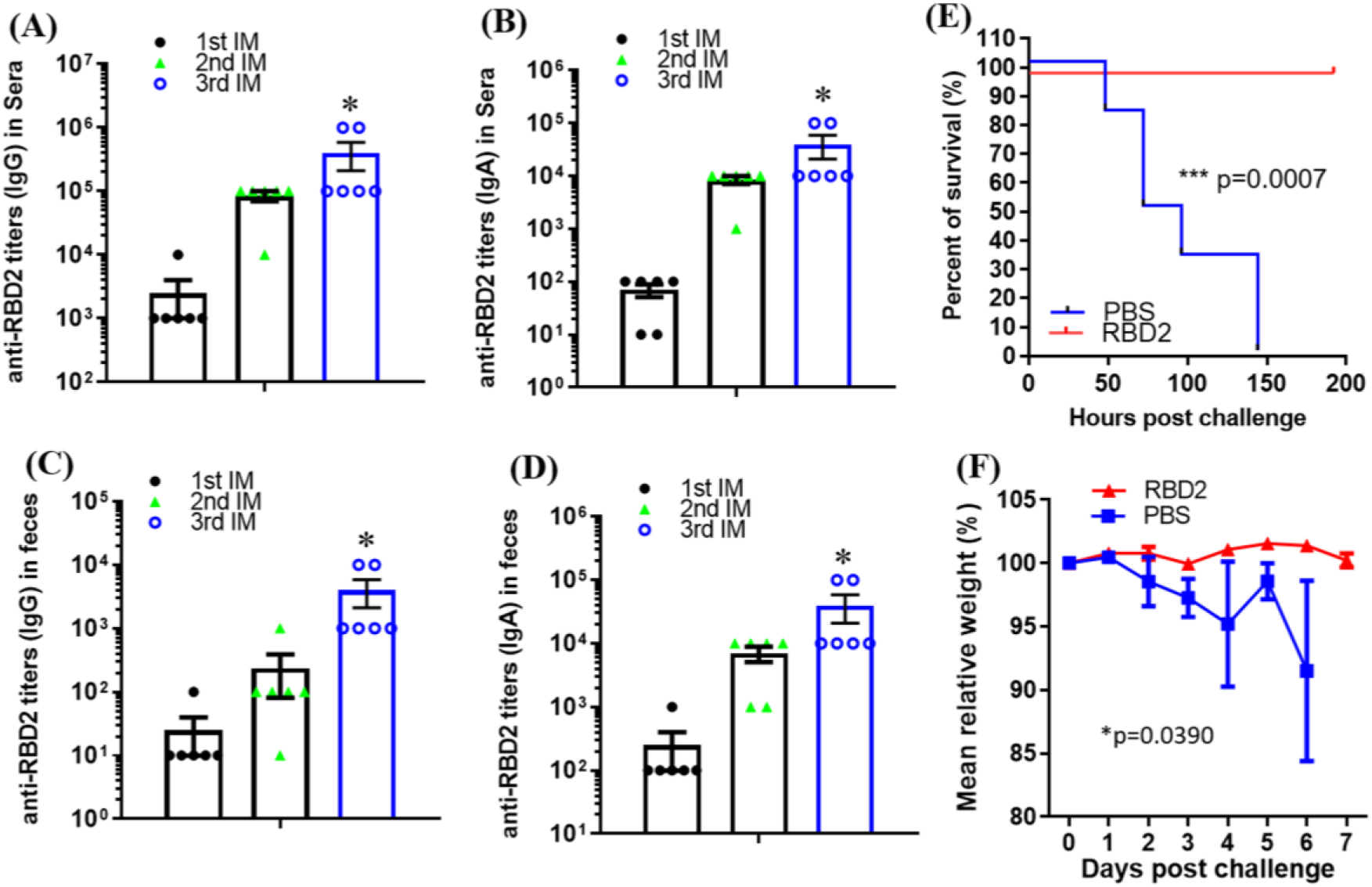
Immunization with RBD2 induces significant anti-RBD2 antibody responses and protects hamsters against infection with *C. difficile* strain DSM 101085. Groups of Golden Syrian hamsters (n = 6 per group) were immunized intramuscularly with 30 µg of RBD2 or PBS (200 µL), formulated with alum, three times at 14-day intervals. Serum and fecal samples were collected after each immunization, and anti-RBD2 IgG and IgA titers were determined by ELISA. **(A–D)** Anti-RBD2 IgG and IgA antibody responses in serum and fecal samples. Two weeks after the third immunization, hamsters received clindamycin intraperitoneally (i.p.; 30 mg/kg/day for 3 days), followed by oral challenge with 2 × 10⁴ spores of *C. difficile* strain DSM 101085 by gavage. **(E)** Survival after challenge. **(F)** Changes in body weight after challenge. Survival differences were analyzed using Kaplan–Meier survival analysis with the log-rank test. The experiment was independently performed twice, and data from one representative experiment are presented as the mean ± SEM. \**p* < 0.05; \*\**p* < 0.001 compared with antibody titers after the first immunization (1st IM).

## Discussion

*C. difficile* binary toxin (CDT) has increasingly been recognized as an important virulence factor in CDI, particularly in epidemic and hypervirulent strains. Although TcdA and TcdB remain the major toxins responsible for CDI, CDT is present in a substantial subset of clinical isolates and has been associated with increased disease severity and recurrence ^6,7,20^. Moreover, clinical isolates producing CDT in the absence of TcdA and TcdB can retain virulence^8,9^, supporting an independent contribution of CDT to disease. Nevertheless, previous clinical vaccine candidates against CDI have primarily focused on TcdA and TcdB^12,21^, leaving CDT largely unaddressed. In the present study, we identified the second receptor-binding domain of CDTb, RBD2, as a highly conserved, immunogenic, and protective antigen and provided evidence that RBD2 represents a promising component for incorporation into a broader toxin-based vaccine against CDI.

A major finding of this study is the marked functional difference between RBD1 and RBD2. Although in silico B cell epitope analysis predicted extensive immunogenic regions in both domains, immunization experiments demonstrated that antigenicity alone did not predict protective activity. RBD1 immunization generated substantial RBD1-specific antibody responses but failed to protect mice against lethal CDT challenge. In contrast, RBD2 and RBD1+2 immunization generated antibodies that recognized intact CDTb and provided complete protection against lethal CDT challenge under the experimental conditions used here. These observations indicate that the location and functional relevance of antibody epitopes within CDTb may be more important for protection than the overall magnitude of domain-specific antibody responses.

The protective activity of RBD2 is consistent with structural studies defining a unique role for this domain in CDTb assembly and toxicity^18^. Unlike RBD1, RBD2 has no counterpart in several other well-characterized binary toxins and forms extensive intermolecular contacts between CDTb heptamers. Structural analyses of activated CDTb identified symmetric and asymmetric di-heptameric assemblies, in which RBD2 contributes critically to the interface connecting the two heptameric units^18^. Deletion of RBD2 markedly reduces CDTb cytotoxicity, further supporting an essential role for this domain in productive toxin function ^18^. Thus, RBD2 represents an unusual structural vulnerability of CDT that may be particularly amenable to antibody-mediated interference.

Our functional experiments provide further support for this interpretation. Addition of recombinant RBD2 or RBD1+2, but not RBD1, significantly inhibited CDT-induced cell rounding. More importantly, sera from animals immunized with RBD2 or RBD1+2 neutralized CDT-mediated cell rounding in a dilution-dependent manner, whereas anti-RBD1 serum showed little detectable neutralizing activity. These findings establish a relationship between RBD2-specific antibodies and toxin neutralization and provide a mechanistic explanation for the protection observed *in vivo*. Antibodies recognizing RBD2 may sterically interfere with one or more steps required for productive CDT intoxication, potentially including CDTb oligomerization, receptor-associated events, CDTa recruitment, membrane insertion, or conformational rearrangements necessary for translocation^14,18^. The present experiments do not distinguish between these possibilities, and defining the precise neutralizing epitopes and molecular mechanism of anti-RBD2 antibodies will be an important subject for future investigation.

Another important property of a vaccine antigen is conservation among circulating strains. Our sequence analyses showed that both CDTb and RBD2 are highly conserved among representative CDT-producing *C. difficile* ribotypes and toxinotypes. Although several amino-acid variations were identified within RBD2, these variations were concentrated primarily toward the N-terminal portion of the domain, whereas the C-terminal region was particularly conserved. Previous epidemiological and genomic studies have demonstrated the broad distribution of CDT among multiple clinically important ribotypes^20,22^. The conservation of RBD2, together with its critical functional role, therefore, supports its potential for providing broad coverage against CDT-producing strains. Broader sequence analyses encompassing larger collections of geographically and temporally diverse clinical isolates will be necessary to establish the extent of RBD2 conservation at the population level.

The protective efficacy of RBD2 was further supported using two complementary animal models. In the direct toxin-challenge model, all non-immunized and RBD1-immunized mice succumbed rapidly after challenge with CDTa plus activated CDTb, whereas all animals immunized with RBD2 or RBD1+2 survived. Activated CDTb alone did not cause mortality, confirming that lethality under these conditions required the functional binary toxin rather than nonspecific toxicity of the CDTb component. Previous studies have established that CDT toxicity depends on the coordinated functions of the enzymatic CDTa component and the binding/translocation CDTb component^13–15^. Importantly, the protective effect of RBD2 was not restricted to administration of purified toxin. RBD2 immunization also significantly protected hamsters against infection with the A^−^B^−^CDT^+^ *C. difficile* strain DSM101085. Because this strain lacks TcdA and TcdB, this model provides an opportunity to examine CDT-directed immunity without confounding toxicity from the two large clostridial toxins.

Our findings also have implications for the design of next-generation CDI vaccines. Previous toxin-based vaccine approaches have predominantly targeted TcdA and TcdB. Such strategies may provide substantial protection against the major manifestations of CDI but do not directly neutralize CDT. This may be particularly relevant for infections caused by CDT-positive strains, including epidemic ribotypes such as RT027 and RT078. Indeed, CDT has been proposed as an additional therapeutic and vaccine target because of its association with clinically important strains and its distinct mechanism of action ^14^. The present findings suggest that RBD2 could serve as a relatively small and defined antigenic component that could be combined with TcdA- and TcdB-derived immunogens to broaden toxin coverage. A multivalent vaccine incorporating protective determinants from TcdA, TcdB, and CDT therefore warrants further evaluation.

Interestingly, inclusion of RBD1 together with RBD2 did not provide an obvious protective advantage over RBD2 alone in the mouse toxin-challenge model. Both RBD2 and RBD1+2 produced complete survival, whereas RBD1 alone was nonprotective. These results suggest that RBD2 contains the principal protective determinants within the C-terminal receptor-binding region of CDTb. From a vaccine development perspective, this observation favors RBD2 as the more streamlined antigen. Nevertheless, the present study was not designed to determine whether RBD1+2 and RBD2 differ quantitatively in neutralizing-antibody potency, durability of immunity, or protection against different CDT-producing strains.

Several limitations should be considered. First, the infection studies employed a CDT-only strain specifically to isolate the contribution of CDT; future studies should evaluate RBD2-containing vaccines against clinically relevant TcdA^+^TcdB^+^CDT^+^ strains, particularly epidemic ribotypes such as RT027 and RT078. Second, although serum neutralization correlated strongly with protection, the specific antibody epitopes and molecular mechanisms responsible for neutralization remain unknown. Structural mapping of protective epitopes and characterization of RBD2-specific monoclonal antibodies could determine whether neutralization results primarily from disruption of CDTb oligomerization, receptor interactions, CDTa association, pore formation, or another step in toxin entry^13,18^.

In summary, this study identifies RBD2 of CDTb as a conserved and functionally important protective antigen against *C. difficile* binary toxin. In contrast to RBD1, RBD2 elicited antibodies capable of neutralizing CDT-mediated cytotoxicity and provided protection against both direct lethal CDT challenge in mice and infection with a CDT-only *C. difficile* strain in hamsters. These functional findings are consistent with structural studies demonstrating the unique contribution of RBD2 to CDTb macromolecular assembly and toxicity. Together, our results establish RBD2 as a promising target for CDT-directed immunity and support its further development as a component of a multivalent vaccine designed to provide protection against all three major *C. difficile* toxins.

## Materials and Methods

### Homology analysis of CDTb and RBD2

CDT^+^ *C. difficile* ribotypes were chosen for analysis based on previous studies ^20,23^. A search was performed in Google Scholar and various genome databases including GenBank (National Center for Biotechnology Information) and the Enterobase *Clostridioides* database ^24^ to identify sequenced *C. difficile* strains from each ribotype (**Table 1**). Enterobase strain ribotypes were specified in each database entry, whereas the ribotypes of other strains were identified using literature sources ^25–31^. while at least two genomes were selected for analysis, with three genomes being used in most cases. CDTb amino acid sequences were mined from each genome before performing a MUSCLE alignment in MegaX software ^32^ on default parameters. Then, a Maximum likelihood phylogenetic trees were constructed in MegaX with 500 bootstrap replicates. The cluster patterns of the phylogenetic trees were used to order the sequences of CdtB or the RBD2 domain of CdtB for a second MUSCLE alignment on the MPI Bioinformatics Toolkit server ^33,34^, as this application produced an output file suitable for visualization using Jalview ^35^. Jalview calculates conservation scores for MUSCLE alignments according to a previously defined algorithm ^36^ that assesses both the amino acid identity as well as physico-chemical properties of the amino acids at a given position to produce a score between zero (no similarities) and eleven (identical amino acids).

### Animals

All animal studies were conducted in accordance with the Guide for the Care and Use of Laboratory Animals of the National Institutes of Health and were approved by the Institutional Animal Care and Use Committee (IACUC) at the University of South Florida. Wild-type C57BL/6 mice and Golden Syrian hamsters were purchased from Charles River Laboratories. Group sizes were selected based on prior experimental data and the anticipated large difference in survival between immunized and non-immunized animals in these lethal challenge models. Assuming 0% survival in non-immunized controls and 90% survival in immunized animals, six animals per group provides approximately 89% power at a two-sided α level of 0.05. Accordingly, six animals per group, balanced by sex (three females and three males), were used for both the mouse CDT challenge and hamster *C. difficile* infection studies.

### Protein expression and purification

Gene sequences encoding the components of CDT (CDTa and CDTb), receptor-binding domain 1 (RBD1), receptor-binding domain 2 (RBD2), or the combined RBD1 and RBD2 domains (RBD1+2) from *C. difficile* R20291 were cloned into pET28a and expressed in *E. coli* BL21(DE3) with an N-terminal His tag. Recombinant proteins were purified from bacterial lysates by Ni-affinity chromatography. CDTb was activated by incubation with trypsin at 0.2 μg of trypsin per μg of protein for 30 min at 37°C. For immunization studies, RBD1, RBD2, and RBD1+RBD2 (RBD1+2) proteins were further purified using Endotoxin Removal Spin Columns (Pierce™) according to the manufacturer’s instructions.

### Mouse immunization and subsequent challenge with CDTa/mCDTb

C57BL/6 mice of both sexes were housed under the same conditions. Food, water, bedding, and cages were autoclaved. Mice (n = 12 per group) were immunized intramuscularly three times at 14-day intervals with 10 µg of purified RBD1, 10 µg of RBD2, or 20 µg of RBD1+2 in phosphate-buffered saline (PBS), with alum as an adjuvant for each immunization. Non-immunized mice (n = 6 per group) served as controls. Serum samples were collected following immunization. Fourteen days after the third immunization, immunized and control mice were challenged intraperitoneally (i.p.) with a lethal dose of CDT consisting of 260 ng of CDTa and 2,870 ng of activated CDTb per mouse (CDTa:mCDTb at a 1:7 molar ratio; n = 6 per group) or with mCDTb alone (2,870 ng/mouse; n = 6 per group). Mice were monitored for survival and clinical signs of disease for 72 h after challenge.

### ELISA for anti-RBD1, anti-RBD2, anti-RBD1+2, anti-CDTb IgG and IgA

ELISA assays were performed as previously described ^37^. Briefly, Costar 96-well ELISA plates were coated with 100 µL/well of RBD1, RBD2, RBD1+2, or CDTb (0.5 µg/mL) at 4°C overnight. After removal of unbound material by washing, the plates were blocked with 300 µL/well of blocking buffer (PBS containing 5% dry milk) for 2 h at room temperature. After washing, 100 µL of serially 10-fold-diluted serum samples were added to each well and incubated for 1.5 h at room temperature. The plates were then washed with PBS, followed by the addition of 100 µL/well of HRP-conjugated anti-mouse IgG (1:3,000) or anti-mouse IgA and incubation for 30 min to 1 h at room temperature. After washing with PBS, TMB substrate was added, and color was allowed to develop for 5–30 min at room temperature. The reaction was stopped by the addition of H₂SO₄, and absorbance at 450 nm (OD₄₅₀) was measured using a microplate reader. Anti-RBD1, anti-RBD2, anti-RBD1+2, or anti-CDTb IgG and IgA endpoint titers were defined as the highest serum dilution at which the OD₄₅₀ was at least twofold greater than that of serum from non-immunized control mice.

### Cell culture and intoxication experiments

Vero cells were used to assess CDT-mediated cytotoxicity. Cells were cultured in EMEM supplemented with 10% fetal bovine serum (FBS) in 48-well plates to approximately 95% confluence (∼5 × 10⁴ cells/well). Cells were then treated with the indicated toxin components and concentrations, as specified in the corresponding figure legends, and incubated at 37°C in a humidified atmosphere containing 5% CO₂. After 2 h of incubation, cell images were acquired using a BZ-X800 microscope (Keyence, IL, USA). The percentage of rounded cells was determined from the acquired images.

### Hamster immunization and hamster model of *C. difficile* infection

Golden Syrian hamsters of both sexes (male = 1:1) were individually housed under the same conditions. Hamsters (n = 6 per group) were immunized intramuscularly with 30 µg of RBD2 formulated with alum as an adjuvant three times at 14-day intervals. Control hamsters (n = 6) received an equivalent volume of PBS. Serum and fecal samples were collected, and anti-RBD2 IgG and IgA titers were determined by ELISA. Two weeks after the third immunization, hamsters received clindamycin intraperitoneally (i.p.; 30 mg/kg/day for 3 days), followed by oral challenge with 2 × 10⁴ spores of *C. difficile* strain DSM 101085 by gavage. Animals were monitored for 7 days after challenge for diarrhea and other clinical signs of CDI.

### Statistical analysis

Animal survival was analyzed using Kaplan–Meier survival analysis with the log-rank test. Comparisons between two groups were performed using an unpaired Student’s t-test. For comparisons involving more than two groups, one-way analysis of variance (ANOVA) followed by Bonferroni’s or Dunnett’s multiple-comparisons test was used, as appropriate and indicated in the figure legends. Data is presented as the mean ± standard error of the mean (SEM). Differences were considered statistically significant at p < 0.05. Statistical significance is indicated as follows: p < 0.05 (*), p < 0.01 (**), p < 0.001 (***), and p < 0.0001(****); ns, not significant. All statistical analyses were performed using GraphPad Prism software.

## Author Contributions

XS conceived, designed, and supervised the project and participated in data analysis. SW and JH performed the experiments and analyzed the data. YN contributed to data analysis. HBK contributed to data analysis and manuscript preparation. All authors contributed to the writing and revision of the manuscript and approved the final version.

## Acknowledgement

The authors thank all members of the Sun Laboratory for their assistance with this project. This work was supported in part by grants from the National Institutes of Health (2R01AI132711, R01AI186760, R01AI149852 and R21AI183094), the USF Center for Antimicrobial Resistance, the USF CREATE Awards, the Anthony Gagliardi Memorial Foundation, and the Florida High Tech Corridor Early-Stage Innovation Award to X.S.

## Data Availability Statement

The data generated and analyzed in this study is available from the authors upon reasonable request.

## Supplemental Figures

**Supplemental Fig. 1.**
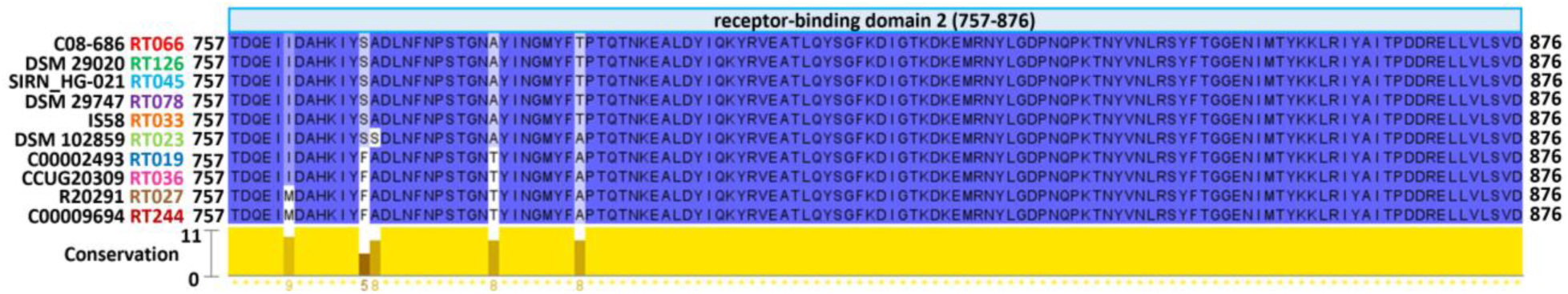
Sequence conservation of RBD2 among representative *C. difficile* strains. MUSCLE alignments of RBD2 amino acid sequences were visualized using Jalview. Conservation scores ranging from 0 (no conservation) to 11 (complete conservation) were calculated for each amino acid position using Jalview (see Methods). The ribotype (RT) of each source strain is indicated adjacent to the strain name and color-coded for ease of identification.

