## Supplementary figures and images for "Receptor-binding domain 2 of *Clostridioides difficile* binary toxin as a promising vaccine component against *C. difficile* infection"

### Supplemental figure 1

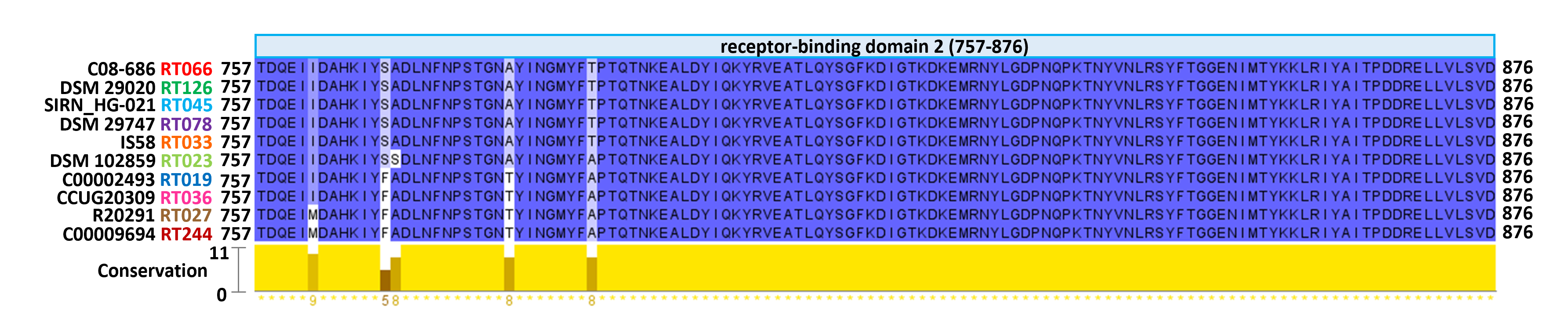
